# Regulating microbiome metabolic stability for stable indigenous liquor fermentation

**DOI:** 10.1101/2023.04.21.537800

**Authors:** Yuwei Tan, Yang Zhu, René H. Wijffels, William T. Scott, Yan Xu, Vitor Martins dos Santos

**Affiliations:** Laboratory of Brewing Microbiology and Applied Enzymology, Key Laboratory of Industrial Biotechnology of Ministry of Education, Jiangnan University, Wuxi, Jiangsu, China; Bioprocess Engineering, Wageningen University and Research, P.O. Box 16, 6700AA Wageningen, Netherlands; Faculty Biosciences and Aquaculture, Nord University, N-8049 Bodø, Norway; Laboratory of Systems and Synthetic Biology, Wageningen University and Research, Stippeneng 4, 6708 WE Wageningen, Netherlands; UNLOCK, Wageningen University & Research and Delft University of Technology, Netherlands

## Abstract

**Background:** Regulating microbial metabolic stability is an ever-challenging goal in the food industry to ensure the productivity and quality of fermented foods. The microbiome underlying traditional Chinese liquor fermentation is such a representative microbiome metabolism that is affected by many dynamic abiotic/biotic factors. The complex microbial activities bring beneficial qualities (complex and rich aroma profiles, *etc.*) to the fermented product, but can also cause unstable fermentation outcomes. Here, we designed a three-step experiment (abiotic regulation; biotic regulation; lab-scale validation) to explore which factors cause unstable fermentation outcomes and how to regulate microbiome metabolic functional stability accordingly.

**Results:** We found that 30.5% industrial fermentation of traditional Chinese liquor outcomes could be precisely predicted by initial abiotic factors. We could ensure the stability of partial fermentation batches by regulating the initial ratio of acidity to reducing sugar, moisture, and starch. Furthermore, in two representative unpredictable fermentation batches (named batch A and batch B), we found that unstable fermentation outcomes occurred even with similar initial abiotic factors after a dynamic three-phase fermentation. Unstable fermentation batches showed fluctuations in microbial community assembly that affected fermentation stability by altering the beneficial distribution (metabolic flux) of redundant metabolic pathways between yeasts and Lactobacilli. The metabolism of batch B was more stable than that of batch A due to the consistent overexpression of a specific set of bacterial metabolic genes. In repeated feed-batch fermentation processes, the difference in metabolic functional stability between the two batches was amplified 9.02 times. Batch B had significantly lower microbiome metabolic fluctuations than batch A, with higher robustness and lower complexity of the metabolic functional network. Moreover, we found that adjusting the initial microbial inoculation ratio could regulate both the metabolic beneficial distribution and temporal metabolic fluctuations of the microbiome to appropriately reduce the instability caused by biotic factors.

**Conclusions:** This study demonstrates that rationally regulating initial parameters and microbial inoculation ratio is a practical strategy to optimize indigenous liquor fermentation. The stable microbial beneficial distribution and high metabolic robustness are essential to obtain the ideal microbiome metabolic stability. Our study provides insights and shows the feasibility of enhancing metabolic functional stability through initial conditions in dynamic microbial ecosystems.

## Introduction

Global demand for microbiome-fermented products (fermented foods, *etc.*) is rapidly increasing [1-3]. Stable microbiome metabolic functionality is essential for the sustainable production of consistent high-quality products [4, 5]. However, improving microbiome metabolic stability is often limited by the fluctuations in physiochemical parameters [6], biodiversity [7], complexity [8, 9], and interaction types [10, 11] of the microbial ecosystem. The effect of these variables can be easily tested in tractable food fermentation ecosystems, and consequently enables optimization to benefit the fermentation control to produce consistent high-quality fermented food and beverages [12].

However, most indigenously fermented foods are produced nowadays by fermentation without scientifically-based control, with the consequence of productivity and quality fluctuations [13]. Tremendous efforts are made to find potential key biotic (biodiversity, *etc.*) and abiotic (physiochemical parameters, *etc.*) factors to ensure a stable fermentation process [14-18]. One fundamental and crucial question is which and how the biotic and/or abiotic factors determine the process stability [15, 19], for the purpose of regulating the fermentation stability by key biotic and abiotic factors. Scientifically-based fermentation control is often achievable in less-complex fermentations (for example pure or defined mixed-culture food fermentation), but difficult to achieve in highly-complex fermentations (for example indigenous solid-state fermentation). In less-complex fermentations, industrial production and lab-scale fermentations provide evidence that temperature, pH, moisture, and lactate are key abiotic process parameters influencing the fermentation stability [20, 21]. Based on these parameters, regulating the fermentation stability is already feasible by statistic models [22]. However, regulating the abiotic parameters is frequently inefficient to ensure a stable high-complexity indigenous fermentation. Sometimes, accidental fermentation fluctuation can still happen when abiotic parameters are kept constant at the start of batches in indigenous solid-state fermentation processes [20, 23, 24]. Such accidental fermentation outcomes are usually ascribed to biotic factors like differences in initial microbial composition and microbial quantity [25, 26]. Nevertheless, how initial biotic factors affect microbial metabolic functional stability remains unclear, which also hampers the efficient prediction and control of high-complexity indigenous fermentations.

Among high-complexity indigenous fermentations, cereal-based indigenous fermentation of Chinese liquor (or called *baijiu* in Chinese) is a unique representative process with saccharification and indigenous fermentation simultaneously and involve complex microbiome of moulds, yeasts and bacteria [27]. Huge data collection of *baijiu* industrial production provides the possibility to trace fermentation batches with recorded or measured biotic and abiotic factors that can affect the quality stability of the end-product. These industrial data can be seen as a rich resource to correlate the parameters with fermentation stability. Generally, quality and productivity fluctuation of end-product among fermentation batches are mainly governed by successive fermentation with yeasts, moulds and bacteria [13]. Here, we aim to achieve more stable *baijiu* fermentation by revealing the associations of abiotic or biotic factors with fermentation stability. Specifically, we target three key unsolved challenges: (i) how to classify unstable fermentation batches caused by abiotic or biotic factors; (ii) how the abiotic or biotic factors contribute to the fermentation stability, and (iii) how to improve fermentation stability by scientifically based control of abiotic or biotic factors at the start of fermentation.

Through integration of statistics, machine learning and multi-omics technologies, we could solve the three challenges in complex fermentation processes through three research steps [28-30]. First, machine learning and prediction modelling can help calculate and classify whether abiotic or biotic factors cause unstable fermentation outcomes [31, 32]. Second, multi-omics measurement can help understand the reason why the key factors contribute to microbiome metabolic functional stability during fermentation processes [33]. Third, creating tools or strategies to regulate microbiome metabolic function stability for stable fermentation would be feasible.

Here, we show how to develop a strategy to achieve more stable *baijiu* fermentation as a representative example, through the three-step experiment (Fig. 1). We first tracked measurements of 6 abiotic parameters in 1009 industrial fermentation batches. Measurement data were mathematically modelled to correlate the 6 parameters with fermentation yield and quality to find the cause of fermentation instability. Then we selected two unpredictable fermentation batches to further reveal how microbial factors other than abiotic parameters caused the fluctuating fermentation. To identify associations of biotic factors with fermentation stability during both short-term and long-term fermentation processes and the underlying mechanism, we utilized a range of techniques, including amplicon sequencing, meta-transcription sequencing, metagenomic sequencing, bioinformatics tools, statistical analysis, and electron microscopy. Finally, to validate the effects of the key factors, we designed simulated experiments with the purpose of regulating the fermentation stability by altering the initial microbial inoculation. We assessed the stability of *in situ* fermentations using the metabolic functional index, and the stability of simulated fermentations using the standard deviation of metabolic profiles. Through substantial analyses, we understand the effect of initial conditions on microbiome metabolic heterogeneity and show how it may help regulate the stability of high-complexity indigenous liquor fermentation.

**Fig. 1.**
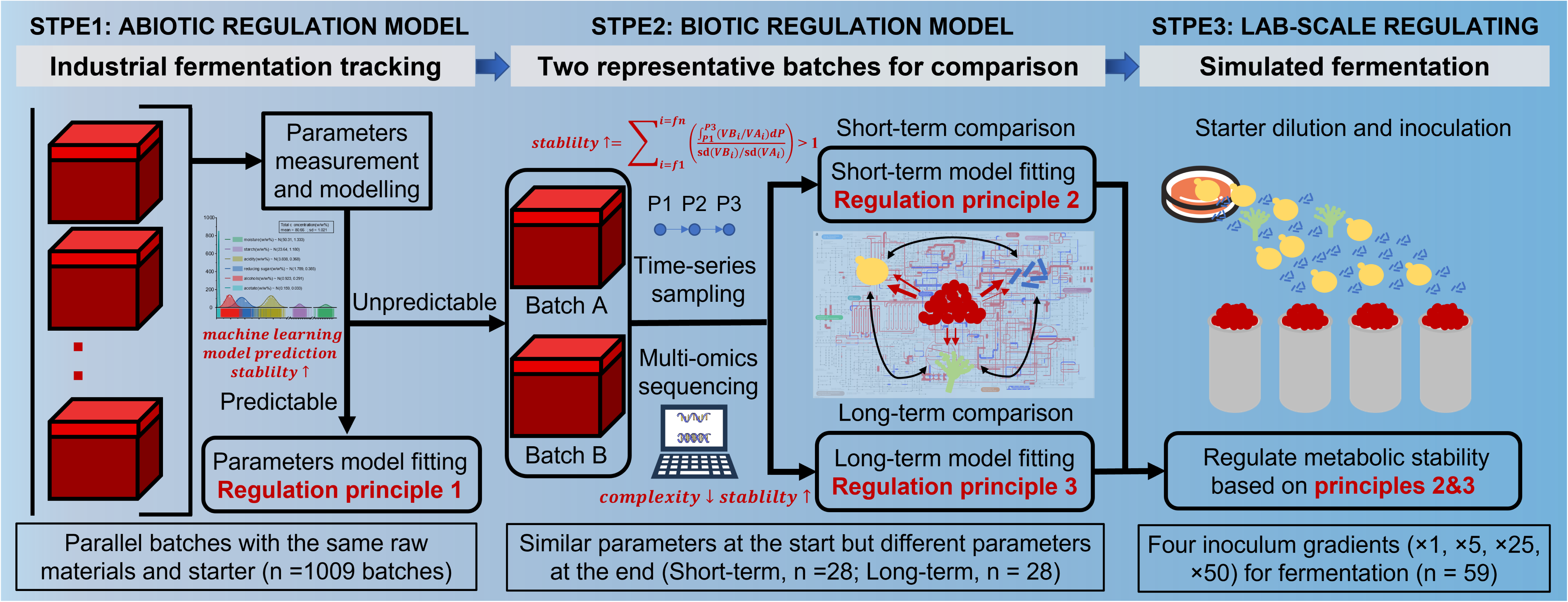
Graphic abstract of three-step experimental design. First, we classified unstable fermentation outcomes caused by initial abiotic factors based on statistic-machine learning combined modelling of 1009 industrial fermentation batches. Second, we revealed the short-term and long-term effects of initial biotic factors on metabolic functional stability based on time-series multi-omics sequencing in two representative fermentation batches. Three regulation principles were summarized to ensure metabolic stability of fermentation microbiome. Third, we conducted simulated fermentations with different initial microbial inoculation conditions to validate feasibility of regulation principles.

## Materials and methods

As mentioned above, we designed our experiments in three separated parts that are described in more detail below (Fig. 1).

### Part 1: Abiotic regulation model

#### Tracking of industrial-scale fermentation

To survey and track the reasons for fluctuating fermentation yield and quality, we collected samples from 1009 batches in industrial-scale fermentation pits (a sort of solid-state fermentation reactor/chamber) in a representative *baijiu* factory (27.85° N; 106.38° E). The fermentation was done in sealed pits for about 30 days. Samples were collected every week from at least 4 positions in a pit and then pooled, transferred to the laboratory within 24 hours in a bucket filled with ice and stored at -20 °C for further analysis.

#### Physiochemical analysis of samples

The moisture of samples was measured by a gravimetric method by drying samples at 105 °C for at least 3 h to a constant weight. The acidity was determined by titration with NaOH (0.1 M), with phenolphthalein as the indicator (endpoint of pH 8.2). Starch, alcohols, acetate and reducing sugar were monitored by the method as previously reported [34].

#### Assumptions and statistical uncertainty modelling of fermentation parameters

Mathematic modelling was conducted to reflect the fluctuating fermentation yield and quality and to find which factors (abiotic parameters or biotic triggered) governed fermentation stability. We assumed that the principal component values calculated from six parameters roughly reflected the quality of fermented products. We fitted Gauss equations to describe fluctuating yield and quality of samples at the end of a batch, as described in Eq. 1:

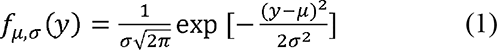

Where y is the dependent variable (related to terminal fermentation parameters), *µ* is the average value, σ is the standard error and π is a constant. The fitting and calculation were conducted in MATLAB (version R2022b). We assumed that the principal component values calculated from six parameters roughly reflected the quality of fermented products. Based on the top two principal component values, we determined the numbers of clusters by machine learning. The number of multiple Gauss models fitting was same as the numbers of clusters. To maximally predict fermentation output by initial parameters, we optimized the model several times as the pipeline shown in Fig. S2b. Partial fermentation batches (26/1009) were deleted as outlier before further modelling. We firstly used linear models to predict fermentation output. However, the sensitivity analysis showed obvious difference between button and top marginal values by Monte Carlo sampling, suggesting potential interactions between input parameters (Table S1). The values of adjusted R square of all linear models were lower than 10, suggesting spline fitting rather than linear fitting would be better for prediction (Fig. S2c). Thus, we established a nonlinear regression model to describe the associations between the value of a given parameter at the start of a batch and at the end of a batch. Taking fermentation yield as a quality marker, we evaluated the model accuracy of yield predictions based on three different thresholds and one classification. The multiple nonlinear regression models are second order models as described in Eq. 2:

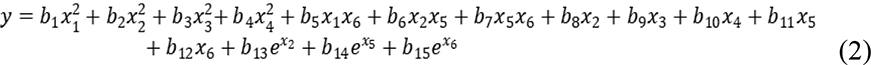

Where *y* is the dependent variable (related to terminal fermentation parameters), *x_1_*through *x_6_* are the initial parameters, *b_1_* through *b_15_* are the regression coefficients, and *e* is a constant. The three different thresholds were alcohols concentration error thresholds of 0.5% (w/w), 0.3% (w/w), and 0.1% (w/w). The classification was three quality types that classified by K-means. Although model could be further optimized by Bayesian inference on the basis of prior distribution as shown in Fig. S2d, the improvement could still be limited without biological insights into the uncertainty.

### Part 2: Biotic regulation model

#### Screening *in situ* representative batches for bioprocess comparison

Using the correlation equations above, physiochemical/abiotic parameters can predict only 30.5% fermentation batches. Thus, other factors (biotic) than abiotic/physiochemical parameters must have played crucial role in determining the productivity and quality. To understand the mechanism behind it, we chose two *in-situ* representative batches according to parameters at the start and the end of batches, named batch A and B. It did not show significant difference in all fermentation parameters of two batches at the start, but showed significant (*P* < 0.05, double-tailed t test) difference in partial fermentation parameters at the end of fermentation. We collected samples every five days during these two fermentation processes. Samples were taken from at least two locations and two depths in a pit and then pooled. Totally, we collected 84 (2 replicates × 7 time points × 2 batches × 3 rounds) samples from batch A and B during the three rounds of fermentation. All samples were transferred to the laboratory within one hour in a bucket filled with ice. Then we conducted fermentation parameters measurement, RNA and DNA sequencing of time-series samples to further reveal the spatiotemporal dynamics of community metabolism.

#### DNA and RNA extraction

Samples were pre-treated with sterile phosphate-buffered saline (0.1 mol/L) and centrifuged at 300 × g for 5 min to obtain the supernatants. Then the supernatants were centrifuged at 11000 × g for 5 min to obtain the sediments.

For DNA extraction, the sediments were cooled and milled in liquid nitrogen and extracted by sodium laurate buffer (sodium laurate 10 g/L, Tris-HCl 0.1 mol/L, NaCl 0.1 mol/L, and EDTA 0.02 mol/L) with phenol: chloroform: isoamyl alcohol (25:24:1) to obtain total DNA. The quality of total DNA was measured by 1% agarose gel electrophoresis and Nano drop 8000 Spectrophotometer (Thermo Scientific, Waltham, MA) (260 nm/280 nm ratio). All genomic DNA of samples were stored at -80°C for further procedure.

For RNA extraction, the sediments were milled with liquid nitrogen and the total RNAs were extracted with sodium laurate buffer (sodium laurate 10 g/L, Tris-HCl 0.1 mol/L, NaCl 0.1 mol/L, EDTA 0.02 mol/L) containing TRIzol (Sigma-Aldrich, St. Louis, MO). Ribo-ZeroTM rRNA Removal Kits (Bacteria) and Ribo-ZeroTM Magnetic Gold Kit (Yeast) (Epicentre, San Diego, CA) were used to remove rRNA from the total RNA. Then, the RNA of samples was stored at -80°C for further procedure.

#### DNA and RNA sequencing

We conducted the DNA amplicon sequencing for 28 samples from the first fermentation round. For the DNA amplicon sequencing, V3-V4 hypervariable region of the 16S rRNA gene and internal transcribed spacer (ITS1/ITS2) region were PCR amplified as previous described [35] and the resulting amplicons were quantified and sequenced on the Illumina Miseq PE300 sequencing platform (Illumina, San Diego, CA).

Based on fermentation phases, we conducted the genomic DNA sequencing only for 28 selected samples (10 samples from the 1^st^ round fermentation; 8 samples from the 2^nd^ round fermentation; 10 samples from the 3^rd^ round fermentation) to evaluate long-term effect of initial biotic factors on functional stability. For the genomic DNA sequencing, the genomic DNA was randomly broken into fragments with a length of about 350 bp a sonicator. Then whole library was prepared through the steps of terminal repair, A-tail addition, sequencing connector, purification, and PCR amplification. After the qualified library pooling, the genomic sequencing was performed on Illumina HiSeq (Illumina, San Diego, CA) platform, which was conducted by the Geneseq Biotechnology Company in Nanjing, China.

Based on fermentation phases, we conducted the RNA sequencing only for 12 selected samples from the first fermentation round to evaluate short-term effect of initial biotic factors on functional stability. For the RNA sequencing, metatranscriptomic libraries were constructed according to the NEBNext® UltraTM RNA Library Prep Kit (Illumina, New England Biolabs, MA) and sequenced on the Illumina Hiseq 2500 platform (Illumina, San Diego, CA), which was conducted by the Allwegene Technology Company in Beijin, China. All sequencing reads can be accessed in the NCBI SRA database.

#### Bioinformatic analysis

For raw DNA sequencing reads, we used Q20 as the quality standard to cut low quality sequences [34]. Then, overlapping reads were merged by fastq-join and primer sequences were removed, only completely assembled reads were utilized for further analysis. The overlap length of merging was 20. The minimum length of fungal sequences was set as 50. The unique sequence set was classified under the threshold of default via Qiime2 (version, 2019-04). Chimeric sequences were identified and removed using Qiime2. The bacterial sequences were mapped to silver132 database for annotation. The fungal sequences were mapped to Unite database for annotation (version 8.2).

For metagenomic DNA sequencing reads, the raw data was filtered based on Q20. Filtered reads were assembled using MEGAHIT (v1.0.6) [36] assembly program (-- min-count 2 --k-min 27 --k-max 87 --k-step 10). Contigs that length less than 500 bp were filtered. Metabat2 was used to do binning process based on contigs. Open reading frame prediction was performed by PRODIGAL [37], and filtered with a length shorter than 100 nt. Using CD-HIT [38] with set (-c 0.95, -G 0, -aS 0.9, -g 1, -d 0) to remove redundancy from the predicted gene sequences. The Bowtie [39] was used to map gene sequences to non-redundant gene catalog with 95% identity. Then the Kyoto Encyclopedia of Genes and Genomes (KEGG) database [40] was used for functional gene annotation at KEGG level 1 and KEGG level 3. We then evaluated the long-term effects of different initial biotic factors on metabolic functional stability based on abundance of assembled genes.

For RNA sequences, we performed species classification analysis, complexity analysis and gene expression abundance analysis. We compared high quality reads to the nonredundant protein database (Nr) and metabolic pathway (KEGG), Gene Ontology (GO), Protein family (Pfam), homologous gene cluster (eggNOG), carbohydrate enzyme (CAZy) to obtain functional annotation information. The sequencing reads for each sample were remapped to the reference sequences using RSEM software [41]. Gene expression levels were measured using the FPKM (Fragments Per Kilobase of transcript per Million fragments) method based on the number of uniquely mapped reads [42]. The DESeq package (ver. 2.1.0) was employed to detect DEGs between two samples [43]. The false discovery rate (FDR) was applied to correct the p-value threshold in multiple tests [44]. An FDR-adjusted p-value (q-value) ≤ 0.05 and a |log2FoldChange| > 1 were used as the threshold to identify significant differences in gene expression in this study. We also evaluated the short-term effects of different initial biotic factors on metabolic functional stability based on gene expression abundance.

#### Calculation of ecological processes

βNTI was calculated to evaluate community assembly patterns, following the protocol described elsewhere [45, 46]. βNTI values that are > −2 or < +2 indicate a stochatic microbial community assembly pattern.

### Part 3: Lab-scale regulating

#### Design of lab-scale simulated solid-state fermentation

To prove the effect of microbial stochastic assembly (dispersal, *etc.*) on the metabolic stability of microbial communities, we designed a lab-scale simulated solid-state fermentation. Multi-species for simulated fermentation were isolated from *Daqu* as shown in Fig. 7a and Fig. S1. The *Daqu* are collected from three different aroma type *baijiu* factories, named Qingxing (QX, means pure aroma), Nongxiang (NX, means strong aroma) and Jiangxiang (JX, means sauce aroma). We first used adequate boiled (sterilized) ddH_2_O to dilute *Daqu* into turbid liquid. Then we centrifuged liquid at 300 × g for 5 min to obtain the supernatants. Then the supernatants were centrifuged at 11000 × g for 5 min to obtain the sediments. The sediments made from three *Daqu* were resuspended by ddH_2_O. All suspensions of isolated species were adjusted to same biomass by sterile water before further dilutions. Then we created 4 dilutions (×1, ×5, ×25 and ×50) of the microbial suspension by sterilized water (details see supplementary Fig. S1) and individually inoculated them into steamed cereals (10%, w/w) to start the fermentation. The control group of each dilution was inoculated with the corresponding microbial solutions without any species via filtering. All fermentations were done under the same conditions (still, 30 °C) for 25 days in 100-ml centrifuge tubes. The fermentations were ended when the weight of whole fermentation tubes changed lower than 0.1 g between two sampling timepoint. Samples were taken and analysed every five days. Samples of group ×1 diluted were collected only three times during the fermentation, because its fermentation ended much earlier than other groups.

#### (HS-SPME-GC-MS) analysis of simulated fermentation samples

To evaluate the fermentation stability, we calculated the Bray-Curtis distance of each sample based on volatile profiles. Volatiles in samples were identified by headspace-solid phase microextraction-gas chromatography-mass spectrometry (HS-SPME-GC-MS) as described above. All samples were pre-treated with 20 mL of sterile saline (1% CaCl_2_, 0.85% NaCl, w/v) in 50 mL centrifuge tubes to collect supernatants after centrifuging at 5000 × g for 10 minutes. Supernatant (8 mL) was added into headspace bottles and mixed with 3 g of NaCl and internal standard (10 µL menthol).

#### (UPLC-MS/MS) analysis of simulated fermentation samples

Metabolites were separated by chromatography on an ExionLCTMAD system (AB Sciex, Framingham, MA) equipped with an ACQUITY UPLC BEH C18 column (100 mm × 2.1 mm i.d., 1.7 µm; Waters, Milford, MA). The mobile phases consisted of 0.1% (w/v) formic acid in water with formic acid (0.1%) (solvent A) and 0.1% formic acid in acetonitrile:isopropanol (1:1, v/v)(solvent B). The sample injection volume was 20 µL and the flow rate was set to 0.4 mL/min. The UPLC system was coupled to a quadrupole-time-of-flight mass spectrometer (Triple TOFTM5600+, AB Sciex, Framingham, MA) equipped with an electrospray ionization source operating in positive mode and negative mode. The detection was done over a mass range from 50 to 1000 m/z.

#### Statistical analysis

All statistical analysis in three experimental parts were conducted as described below. Dynamics of physical and chemical factors were fitted with OriginPro2019. Principal component analysis (PCA), significant tests, microbial correlations, canonical-correlation analysis, random forest model and variance partitioning analysis were conducted by vegan package in R (http://vegan.r-forge.r-project.org/). We calculated the structure equation model to clarify the causality and quantify the effects of microbial stochastic assembly to metabolic functions by IBM SPSS Amos 25. *P*-values were adjusted for nonparametric analysis by the Statistical Package for Social Science (SPSS, version 22). The metabolic functional stability index of microbiome was calculated based on average variation degree [4]. Specifically, we calculated the functional stability by taking the ratio of the mean to standard deviation for various functional categories (KEGG level 1) and individual functional profiles (KEGG level 3) across replicated fermentation samples [47]. As for main metabolic functional categories of two selected batches, we derived a function to determine the change of overall metabolic functional stability between fermentation batches as described in Eq. 3: 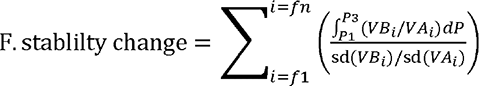 (3). Where *f1, f2…fn* represents different metabolic functional genes. P1, P3 represents fermentation phases. VBi or VAi represents the FPKM values of gene i. Sd(VBi) or sd(VAi) represents standard deviation of the FPKM values for gene i.

#### Data visualization

Data plotting were carried out in Graphpad prism (version 7), OriginPro2021, Microsoft® Excel and Adobe Illustrator CS6. Different gene expressions of overall metabolic functionality mapped to the KEGG metabolic network using iPATH3 [48].

## Results

### Unstable fermentation batches caused by abiotic factors

Fig. 2 shows the fluctuating yield (alcohols concentration) and quality (reflected by six parameters) in industrial-scale fermentation. Total normalized concentration of the 6 parameters was 80.66 ± 1.021% (w/w) at the start of fermentation and 83.44 ± 1.112% (w/w) at the end of fermentation. The six physiochemical parameters all showed normal distribution at the start of batches. The mean value of moisture was 50.31% (w/w), starch was 23.64% (w/w), acidity was 3.838% (w/w), reducing sugar was 1.789% (w/w), alcohols was 0.923% (w/w), and acetate was 0.159% (w/w). At the end of batches, the starch and reducing sugar turned to lognormal distributions, suggesting the depletion of available carbon sources in some batches (Fig. 2a). The unstable physiochemical parameters at the end of batches showed three peaks by Gauss model-based principal component analysis (Fig. 2b). According to fusion levels of top two principal components, we classified unstable fermentation outputs into three quality types (high, medium, and low quality) by K-means (Fig. 2c). The alcohol concentration range of high quality was from 2.43 to 3.46%, medium quality was from 2.19 to 2.42% and low quality was from 0.42 to 2.18%. There were 377 batches of high quality, 295 batches of medium quality and 337 batches of low quality (Dataset 1).

**Fig. 2.**
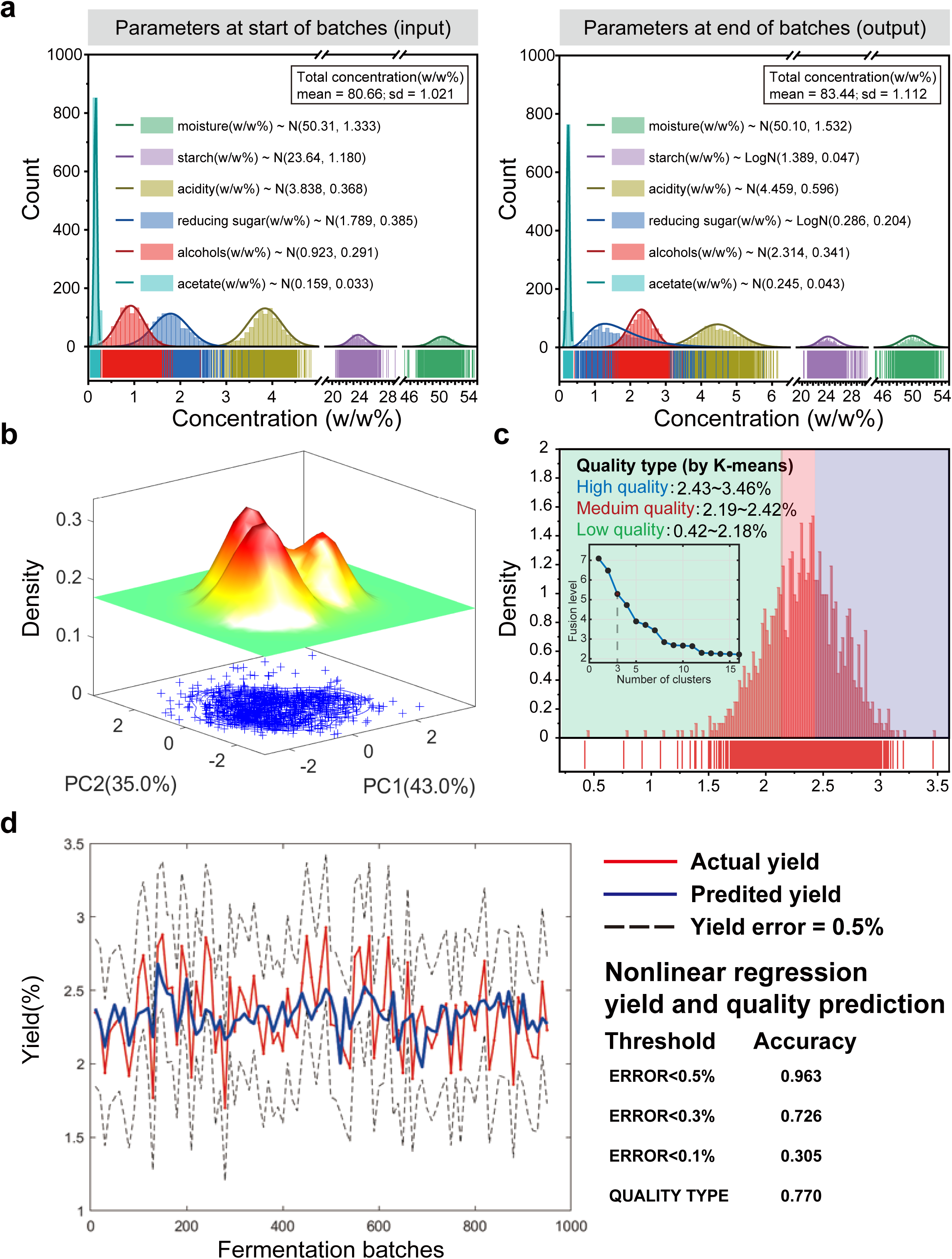
Fermentation outcomes prediction based on initial parameters. (a) Distribution of input and output parameters. (b) Unstable parameters at the end of 1009 batches clustered by multiple gaussians model-based principal component analysis. (c) Unstable batches are classified into three quality types by K-means. The number of both gauss models and quality types was determined by fusion level. (d) Accuracy of yield (%, w/w) prediction by nonlinear regression model.

To predict fermentation outputs by initial input parameters, we conducted statistical uncertainty modelling (Fig. S2a). We found that 30.5% fermentation batches could be precisely (at 0.1% yield error threshold) predicted based on abiotic parameters at the start of batches (Fig. 2d). The ratio of initial acidity to initial reducing sugar concentration (0.182 ≤ ratio ≤ 0.216), initial moisture (2.539 ≤ ratio ≤ 3.296) and initial starch concentration (0.079 ≤ ratio ≤ 0.085) were three key parameters to regulate fermentation stability. As the prediction accuracy increased, the prediction precision dropped rapidly. The prediction accuracy was 72.6% at 0.3% (w/w) yield error threshold and increased to 96.3% at 0.5% (w/w) yield error threshold. The large uncertain precision of yield (0.3% - 0.5%, w/w) suggested that most fluctuating yield cannot be predicted barely based on physiochemical parameters at the start of batches. Nevertheless, the prediction accuracy of fermentation quality types was 77%, which was acceptable.

### Representative two unstable fermentation batches caused by biotic factors

During dynamic *baijiu* fermentation process, physiochemical parameters changed with fermentation time. We divided fermentation process into three phases based on dynamics of physiochemical parameters. Fig. 3a shows the three phases during fermentation process, that is, phase I (0-5 days), phase II (5-21days), and phase III (21-30 days). Fig. 3b shows the dynamics of parameters (normalized by same sample weight) throughout the fermentations. We found that alcohols were mainly generated in phase I and phase II, whereas acids were mainly generated in phase III. Alcohols rapidly increased from 0.272 ± 0.092% (w/w) to 0.582 ± 0.101% in phase I and increased from 0.582 ± 0.101% (w/w) to 0.700 ± 0.086 % in phase II. Acetate (from 0.475 ± 0.064% to 0.613 ± 0.105%, w/w) and acidity (from 0.465 ± 0.067% to 0.579 ± 0.124%, w/w) continuously increased during phase II and phase III. Moisture gently increased during phase I and phase II (from 0.521 ± 0.069% to 0.678 ± 0.071 %, w/w), and decreased afterwards. Starch showed the opposite changing trend to moisture, it gently decreased during phase I and phase II (from 0.459 ± 0.059% to 0.219 ± 0.072 %, w/w). Reducing sugar decreased throughout the fermentation process (Dataset 2). Based on our predicting model, unpredictable batches could be screened by comparing physiochemical parameters in each fermentation phase.

**Fig. 3.**
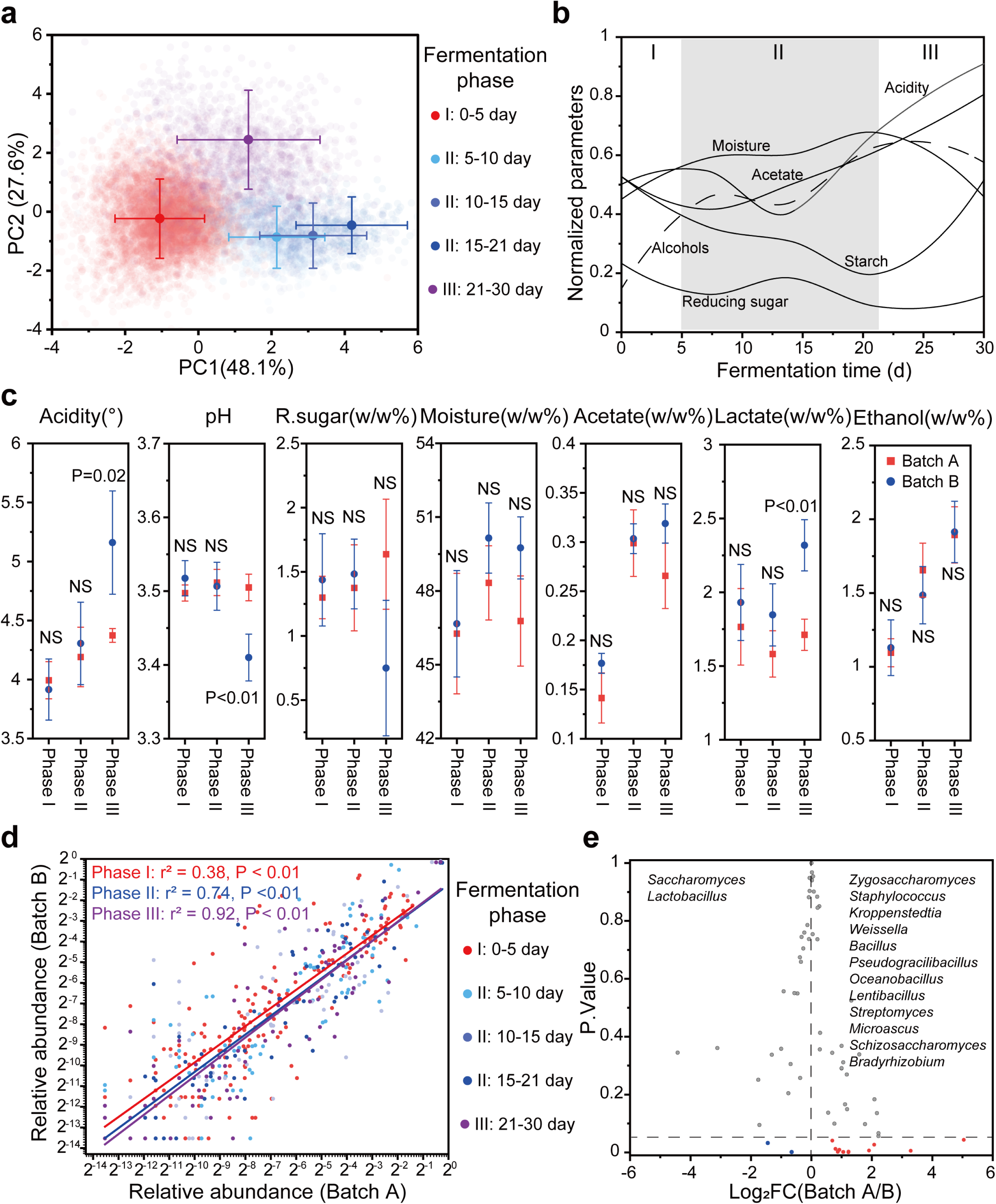
Three-phases dynamic fermentation process. (a) Principal component analysis of parameters showing changes during fermentation. The samples are colored according to the three fermentation phases as determined by parameter clusters. (b) Dynamics of fermentation parameters (acidity, reducing sugar, moisture, acetate, starch, alcohols) during 1009 industrial fermentation batches. All parameters are normalized by sample weights. (c) Fermentation parameters difference (double-tailed t test) between batch A and batch B in each fermentation phase. (d) The relative abundances of microbial genera in batch A are in stark contrast with those in batch B in fermentation phase I but are similar in fermentation phase II and fermentation phase III. Each point represents a microbial genus. (e) Microbial genera with significant (one-way ANOVA test) relative abundance differences.

We selected two representative unstable fermentation batches (named batch A and batch B) after comparison of their parameters. We found all parameters of these two batches were similar in fermentation phase I and II but partial parameters were significantly different in fermentation phase III (Fig. 3c). Acidity, pH and lactate showed significant (*P* < 0.05, tested by one-way ANOVA) difference in phase III. We found batch B had higher acidity, lactate and lower pH than that of batch A. Obviously, the difference of microbial variability appeared earlier than that of physiochemical parameters. Fig. 3d shows that microbial structure of batch A and B was unsimilar in phase I, whereas became more and more similar with fermentation time. Microbial community showed a convergent succession process across fermentation phases. In phase I, the r^2^ of co-correlation was 0.38, then increased to 0.74 in phase II, and finally increased to 0.92 in phase III. Fig. S3 shows two fermentation batches encompassed same 12-13 stably coexisted dominate (average relative abundance > 1%) fungal and bacterial genera but with distinct relative abundances and succession. As shown in Fig. 3e, 12 microbial genera were more abundant in batch A than in batch B, including *Oceanobacillus*, *Bacillus*, *Pseudogracilibacillus*, *Kroppenstedtia*, *Lentibacillus*, *Schizosaccharomyces*, *Streptomyces*, *Weissella*, *Staphylococcus*, *Microascus*, *Zygosaccharomyces* and *Bradyrhizobium*. The relative abundance of *Lactobacillus* and *Saccharomyces* was more abundant in batch B. In addition, Fig. S4 shows that biotic factors of both batch A and B contributed more than abiotic parameters to eight quality related parameters through random forest modelling. Biotic factors contributed to ethanol most (∼80%IncMSE) in batch A and contributed to acetic acid most in batch B (∼75%IncMSE). Abiotic factors contributed to lactate most (∼10%IncMSE) in batch A and contributed to moisture most in batch B (∼10%IncMSE). Thus, we confirmed that these two fermentation batches were representative unstable fermentation batches caused by biotic factors.

### Microbial community assembly patterns of two representative batches

Fig. 4a shows the transmission electron microscopy of pooled fermented samples. The hyphae, yeasts and bacteria were embedded in the grain matrix. We found that spatial distribution and density of microbial cells would change with fermentation phases. Yeasts showed high density in phase II, whereas bacteria showed high density in phase III, suggesting variable dominate species and dynamic community assembly patterns.

**Fig. 4.**
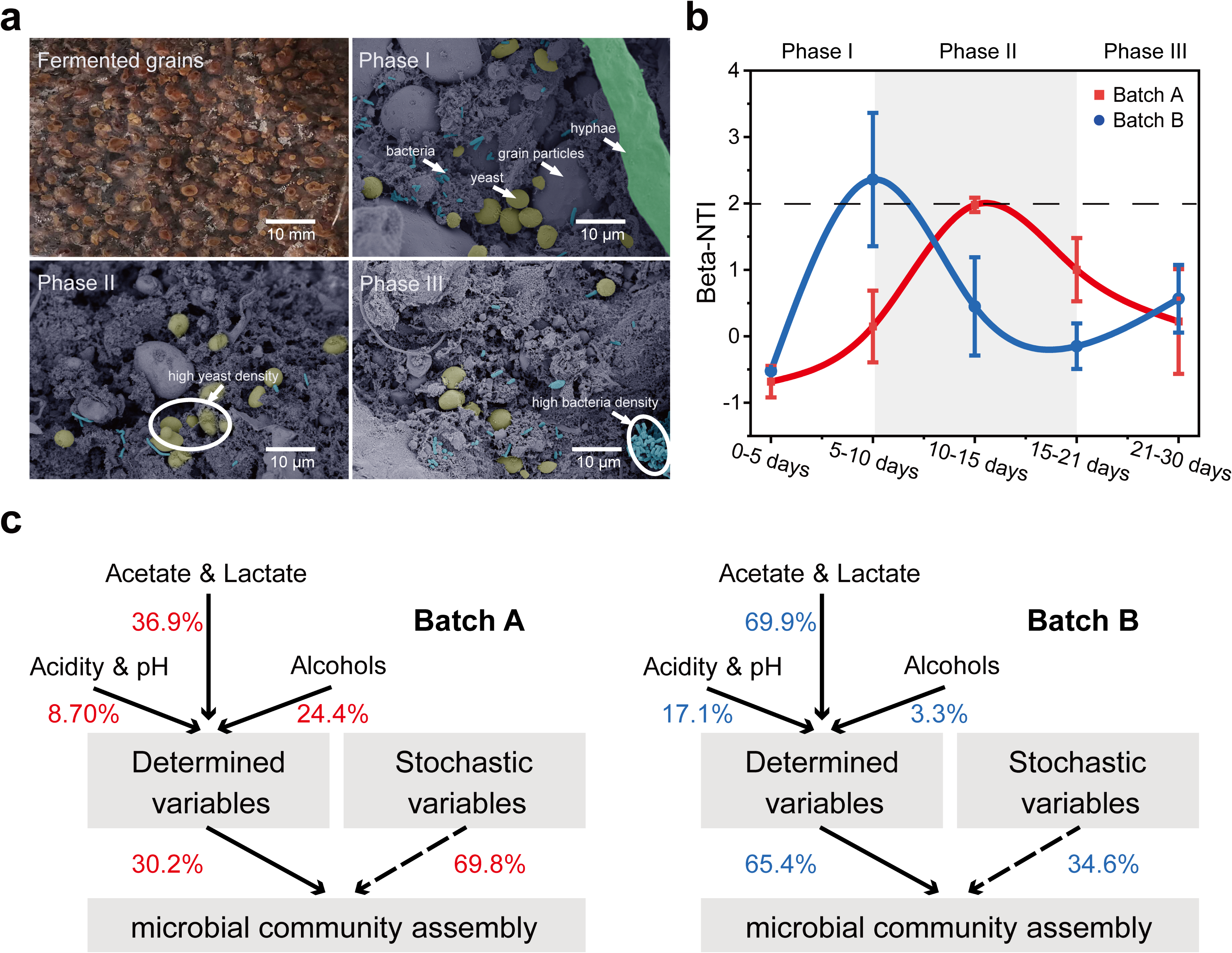
Dynamic traits of microbial assembly. (a) Example image of macroscopic fermented grains (top left), and transmission electron microscopy of hyphae (marked with green), yeasts (marked with yellow), bacteria (marked with blue) embedded in the grain matrix at fermentation phase I (top right), phase II (bottom left) and phase III (bottom right). (b) Community assembly patterns during fermentation process. (c) Variance partitioning analysis of the relative contributions of determined (alcohols, acids, *etc.*) and stochastic variables to variation in microbial community assembly. The solid lines indicate the contribution of the variables. The dashed lines represent the assumed contribution of stochastic variables (1 – determined contributions%).

Fig. 4b shows microbial community assembly was governed by stochastic assembly process during phase I and phase III, whereas the community assembly of phase II was a determined assembly process. Although two representative batches underwent same three-phases assembly process, i.e., a stochastic-determined-stochastic process, the timepoint of phase shift was different. Batch B showed earlier timepoint of phase shift and stronger environment selection than that of batch A during the fermentation. The assembly shifting of batch A happened in the second period of phase II (10-15 days after the start of fermentation), whereas batch B happened in the first period of phase II (5-10 days after the start of fermentation). The maximum βNTI value was about 2.36 ± 1.00 in batch B, whereas the the maximum βNTI value was about 1.98 ± 0.11 in batch A.

We evaluated the relative contributions of determined variables (alcohols, pH, acidity, acids, lactate and acetate) and stochastic variables to community assembly (Fig. 4c). The most important contributors of determined variables in batch A and B were same but with different strength. Specifically, lactate and acetate were the most important abiotic contributors of community assembly in batch A, followed by alcohols and then pH, acidity. Lactate and acetate were also the most important abiotic contributors to community assembly in batch B, but followed by pH, acidity and then alcohols. Meanwhile, we found that the total contributions of these determined factors were 30.2% in batch A and 65.4% in batch B, suggesting the greater importance of microbial stochastic assembly in batch A than that in batch B. Collectively, unstable fermentation caused by initial biotic factors showed fluctuations in community assembly patterns and microbial responses to fermentation parameters.

### Short-term effect of biotic factors on metabolic functional stability

To get deeper insights into community assembly patterns affecting microbial community functions and metabolic stability, we analyzed the dynamics of metabolism-related gene transcriptions. The metabolic functions of the two batches were mainly involved in functional categories of carbon, nitrogen, and sulfur metabolism. According to the FPKM (Fragments Per Kilobase of transcript per Million mapped reads) value of every functional category, we found that the main community metabolic functions changed with the fermentation phases (Fig. 5a). During phase I, the main community functions were “sulfur metabolism”, “terpenoid and polyketide metabolism”, “glyoxylate and dicarboxylate metabolism”, “lipid metabolism” and “TCA cycle”. During phase II and phase III, the main community functions changed to “carbon metabolism”, “butanoate metabolism”, “propanoate metabolism”, “methane metabolism”, “galactose metabolism”, “nitrogen metabolism” and “amino acid metabolism”. The fluctuate metabolic functions between batch A and batch B were reflected in 14 functional categories as shown in Fig. 5a. Most (12/14) microbial metabolic functional categories of batch B were more active than batch A in phase II and phase III according to FPKM values. According to Person’s correlations between βNTI values and FPKM, we found functional categories of “pyruvate metabolism”, “metabolism of xenobiotics” and “glycolysis/gluconeogenesis” were significantly negative correlated with |βNTI| values (Fig. 5b). Based on structure equation model, we further illustrated the causality between metabolic functional expressions and community assembly patterns (Fig. 5c). We assumed that initial biotic factors caused changes in microbiome metabolism between fermentation batches through community assembly. The results of the data fitting did not significantly deny the model assumptions (*P* = 0.343). Different community assembly patterns explained 89.3% fluctuation of “glycolysis”, 42.4% fluctuation of “pyruvate metabolism”, and 42.1% fluctuation of “metabolism of xenobiotics” between two unstable fermentation batches.

**Fig. 5.**
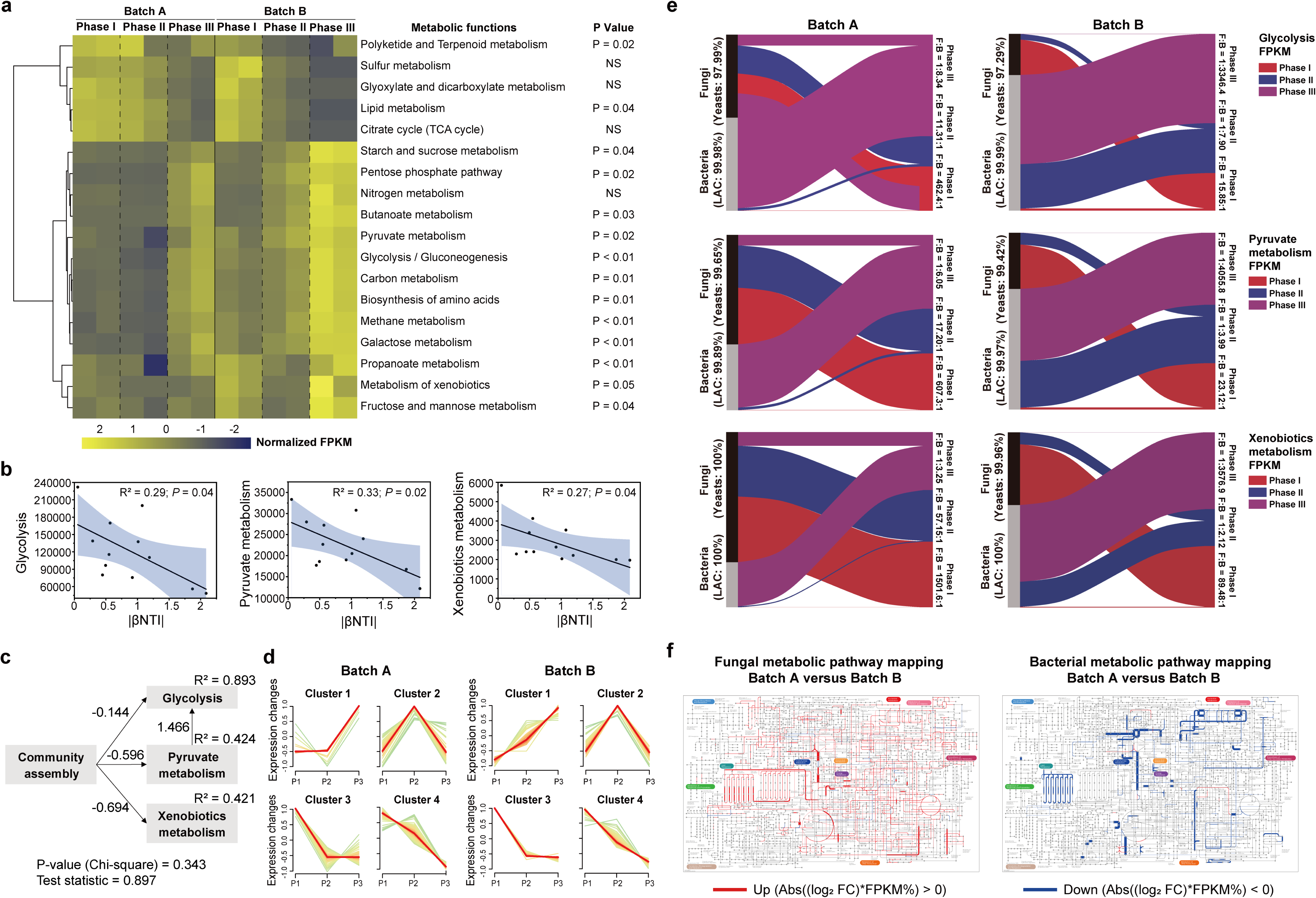
Fermentation instability caused by dynamic microbial beneficial distribution. (a) Significant (double-tailed t test) different transcription of 18 major metabolic functional categories. (b) Significant correlations between metabolic functional categories and community assembly (double tailed t test). (c) Structural equation model indicates significant effects of assembly patterns on three metabolic functional categories. (d) Dynamic gene expression levels among fermentation phases according to FPKM values. (e) Alternative beneficial distribution between yeasts and lactic acid bacteria in three metabolic functional categories. (f) Overall fungal and bacterial metabolic fold change (batch A/batch B) between two batches. The thickness of pathway lines is calculated by Abs ((log_2_ fold change) × average FPKM% values). Red lines represent up-regulated pathways, whereas blue line represent down-regulated genes (Details see dataset 5).

Fig. 5d shows the dynamics of all assembled gene transcriptions within these three function categories in each fermentation phase. There were four clusters of genes in both batch A and batch B according to different expression dynamics, that is, cluster 1 (increased and increased), cluster 2 (increased then decreased), cluster 3 (decreased then stable), and cluster 4 (decreased and decreased). Such dynamics suggested that multiple alternative genes could perform the same metabolic function. For functional categories of “pyruvate metabolism”, “xenobiotics metabolism” and “glycolysis”, we found lactic acid bacteria occupied more than 99% genes transcription in bacteria, whereas yeasts occupied more than 97% genes transcription in fungi. The black-grey bars showed that yeasts were more active in batch A, whereas lactic acid bacteria were more active in batch B (Fig. 5e). For gene expressions of glycolysis in phase I, the cumulated FPKM value of fungi was 462.1 times bigger than that of bacteria in batch A, whereas the ratio was only 15.85 in batch B. Such gene expression difference was similar in pyruvate and xenobiotics metabolism. In addition, we found different turnover time of fungi-bacteria alternative interactions between two batches. Batch B showed turnover of fungi-bacteria alternative metabolism in phase II. Batch A showed such a turnover in phase III. Partial detailed different in alternative fungi-bacteria metabolism flux between the two batches was summarized Fig. S5. We found that the main consumer of substrate in “Pyruvate metabolism” and “Glycolysis” changed between the two batches. *Lactobacillus* was the main consumer of substrate in batch B, whereas *Zygosaccharomyces* was the main consumer of substrate in batch A. In result, different metabolic fluxes between fungi (mainly yeasts) and bacteria (mainly lactic acid bacteria) further caused the instability of overall metabolic functions (Fig. 5f and dataset 3). We found the obvious bacterial overexpression of most genes in batch B, whereas fungal overexpression of most genes in batch A (Fig. 5f). In addition, bacterial gene overexpression focuses on fewer metabolic pathways than fungal gene overexpression. Based on the temporal stability calculated by Eq. 3, we found that the metabolic stability of microbiome was 2% higher in batch B than in batch A for the 18 main functional categories.

### Long-term effect of biotic factors on metabolic functional stability

Fig. 6a shows the fluctuations in biodiversity and functional gene abundance during long-term fermentation processes. During long-term fermentation processes, we found the overall microbial functionality became more and more unstable (Fig. 6b). Compared with the functional stability index of Round 1, the index decreased 1.33% in fermentation Round 2. The decreasing percentage of functional stability index increased to 9.02% in fermentation Round 3. Such constantly increasing functional instability suggested the diffidence in overall functional stability between the two representative batches was amplified. In addition, we found the stability of main metabolic functions in batch B was less affected than in batch A. The metabolic functional stability index of batch B was significant (*P* < 0.05, student’s t-test) higher than that of batch A (Fig. 6c).

**Fig. 6.**
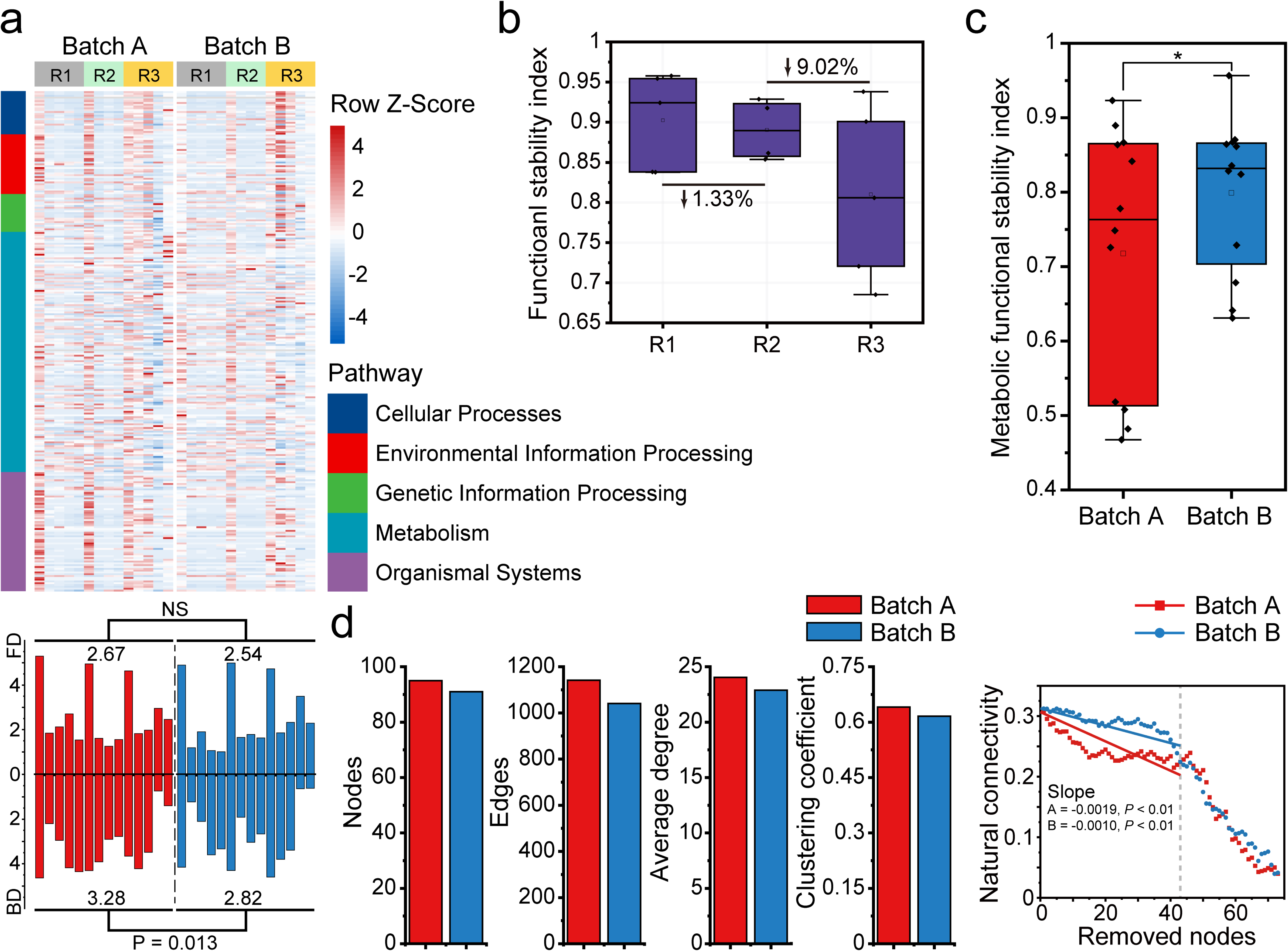
Fermentation instability caused by fluctuating metabolic robustness. (a) Fluctuating functional traits (top), fungal (FD) and bacterial (BD) Shannon diversity (button) of two batches during three-round repeated feed-batch fermentation process. (b) Overall functional stability index among three fermentation rounds. (c) Metabolic functional stability index of batch A and batch B (*, *P* < 0.05, t test). (d) Metabolic network topological properties between batch A and batch B. Robustness linear fitting (one-way ANOVA test) are cut off at 43 removed nodes.

We then compared the topology parameters of metabolic functional networks in batch A and B over the process of the long-term fermentation (Fig. 6d). Batch A showed higher values in nodes, edges, average degree of nodes and clustering coefficient than in batch B, suggesting higher network complexity in batch A. Interestingly, batch B showed better robustness but lower complexity of the metabolic functional network. When removing less than 43 nodes from the metabolic network, we found the resistance of batch A metabolic network was weaker than batch B. When removing more than 43 nodes from the metabolic network, batch A and batch B showed comparable network robustness.

### Optimize fermentation stability by regulating initial microbial inoculation

To improve fermentation stability through initial biotic factors, we designed four inoculation ratios (conducted by dilutions) as an approach to regulate fermentation stability (Fig. S1). Fig. 7a shows species for fermentation were isolated from three different starters (named QX, NX, JX). We found the number and the detailed express level of metabolites were affected through inoculation dilution (Fig. 7b). Dataset 4 shows that the express level of metabolites related to “pyruvate metabolism” and “glycolysis/gluconeogenesis” fluctuated among four inoculation ratios, mainly including citric acid, maltose, lactose, phenylacetic acid, and phenylalanine. The different express level of metabolites evidenced the unstable metabolic flux of microbial community.

**Fig. 7.**
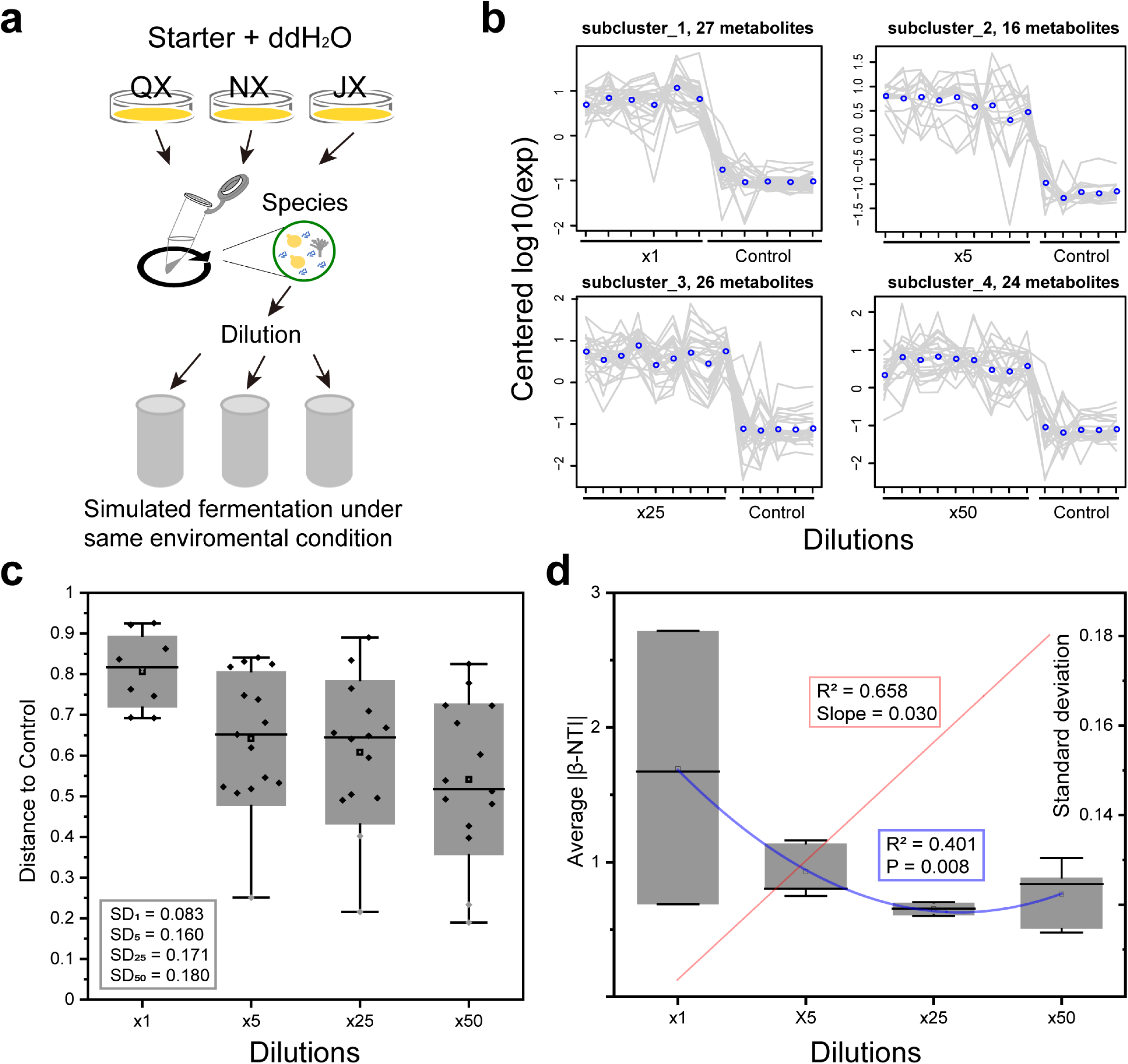
Initial microbial inoculation ratio (by dilutions) affects metabolism fluctuations and community assembly within a simulated fermentation ecosystem. (a) Schematic of the experimental design under different initial inoculation conditions. (b) Changes in metabolites during initial and medium fermentation phases reveal different patterns of metabolic expression levels. The abscissa is each comparative sample group, and the ordinate is the expression level of metabolites in groups. Each grey line represents a metabolite, and the blue line represents the average expression of all metabolites in the sub-cluster. Different metabolites detected by UPLC-MS/MS are grouped into sub-clusters via H cluster. (c) Bray-Curtis distance between the four groups to the control group based on volatile metabolic profiles during fermentation. SD represents the standard deviation of Bray-Curtis distances to reflect metabolic stability. (d) Community assembly (blue) patterns and metabolic stability (red) at four dilution gradients during the first 15 days of fermentation. The intersection of the fitted lines suggests the ideal microbial inoculum ratio.

Fig. S6 shows that the volatile profiles were significantly influenced by inoculation dilution despite the difference in starters. Based on linear fitting results between PC1 values and different dilution degrees, we found that fermentation of the JX group was the most sensitive group to dilution conditions (R^2^ = 0.604, P = 0.00015), whereas the QX group was the most insensitive group (R^2^ = 0.258, P = 0.022). Fig. 7c shows that fermentation with low dilution tended to have high efficiency of raw material conversion. Besides, groups with lower dilution showed higher fermentation stability. For the group with ×1 dilution, the standard deviation of distances was 0.083. For the ×5, ×25 and ×50 dilutions, the standard deviations of distance were 0.160, 0.171 and 0.180, respectively.

Fig. 7d shows that the dilution gradient was linearly related to fermentation stability (R^2^ = 0.658, Slope = 0.030) and second order linearly related to the community assembly pattern (R^2^ = 0.401, *P* = 0.008). Compared with other groups, the group ×5 showed both acceptable community assembly pattern and fermentation stability. The volatile profile of group ×1 was the most stable group but ended fermentation 10 days earlier than other groups due to the strong determined community assembly pattern.

## Discussion

Stable indigenous food fermentation is essential for food productivity, quality and safety as well as sustainability [23, 27]. The more complex a fermentation microbial ecosystem is, the more difficult to achieve a stable fermentation outcome due to dynamic fermentation fluctuations in time, space and composition [49]. In our study, we demonstrated that optimization of high-complexity indigenous liquor fermentation was feasible by regulating the metabolic functional stability of microbiome through initial abiotic and biotic factors. Specifically, regulating initial fermentation parameters would help reduce fermentation instability that caused by abiotic factors. Regulating initial microbial inoculation ratio would help reduce fermentation instability caused by biotic factors. This is supported by the findings of our three-step experiment. Here, we further discussed the implications of our findings and summarized three principles for regulating microbiome metabolic functional stability in similar or less complex microbial ecosystems.

Artificial intelligence (AI) methods and quantitative regression model are effective tools to predict and ensure stable fermentation outcomes [50, 51]. Evaluating model input and output are important to define and classify different productivity, safety and quality types for further model prediction and control [52]. However, how many productivity, safety and quality types should be identified as a minimum to efficiently distinguish each other for further modelling is usually hard to decide. In part one of our experiment, taking six parameters as a simple example, we showed that machine learning-based unsupervised analysis and clustering could help find the most suitable number of different fermentation types (Fig. 2b, 2c). For cases with more parameters (i.e. sensory evaluation), machine learning or deeper machine learning (i.e. neural network) analysis has the potential to help find the best classification for fermentation stability prediction and control [53].

Based on the statistic model combined with machine learning here, our findings demonstrated that, when input fermentation parameters have significant correlations, regulating individual parameters contributed little to fermentation outcomes (Fig. S2). Many input fermentation parameters (acidity, pH, reducing sugar, *etc.*) are parallel driving factors that respond simultaneously to microbial succession at different fermentation phases [34, 54, 55]. The parameters usually exhibit non-linear interactions with one another during dynamic liquor fermentation process [56]. In non-linear microbial ecosystems, the phases shifts of the microbiome are determined by the "energy landscape" [57]. This explained why thousands of industrial liquor fermentation batches exhibited similar three-phase fermentation patterns despite fluctuations in parameters (Fig. 3a, 3b). In our opinion, the essence of unstable liquor fermentation batches was the fluctuation of fermentation parameters around the multiple attractors of the non-linear ecosystem. Such fluctuation (e.g., butterfly effect) is usually affected by initial conditions [58, 59]. Notably, regulating individual initial parameters could enhance stability of one fermentation phase but might have a limited influence on overall fermentation stability. Collectively, our findings from part one of this study supported the first regulation principle, that is, in microbial ecosystems with multiple driving parameters, rationally regulating initial ratio of key parameters could help efficiently reduce metabolic instability that caused by abiotic factors. For metabolic instability caused by biotic factors, unstable fermentation outcomes may occur for a variety of reasons at different fermentation phases. For example, microbial inhibition by fermented products or physiochemical parameters often happens in meddle or later fermentation phases [60], and presence of microorganisms that compete for substrate consumption often happens in initial phase [26, 61]. Here, we defined the three phases of *baijiu* fermentation marked by dynamic physiochemical parameters. Across the three fermentation phases, the *baijiu* microbiome showed a convergent succession process (Fig. 3). The convergent succession in solid-state media is heavily influenced by the initial transcription of microbial genes [26, 34, 62]. In addition, this transcription can vary depending on the microbial immigration and probabilistic spatial dispersal that occur at the beginning of solid-state indigenous fermentation [63]. Fast dispersal rate of species would benefit uptake speed of resources [63-65]. Thus, the convergent succession process is not only reminiscent of microbial spatial exploration theory [64] but also contains elements of consumer-resource models applied to microbial coexistence [66]. At the start of two selected fermentation batches, our results showed that the different major consumers of carbon and nitrogen sources (fungi or bacteria) induced gene expression changes, suggesting regulating initial biotic factors can affect microbial succession and thus metabolism (Fig. 5 and dataset 5). The underlying mechanism is different resources consumers can affect microbial beneficial distribution (metabolic flux) in microbial ecosystems with metabolic division of labour. The benefits distribution of microbial members is essential to metabolic stability [67]. Our short-term fermentation results demonstrated that batches with stable overexpression of bacterial metabolic genes (longer maintenance of stable beneficial distribution) showed superior temporal metabolic functional stability. For instance, higher stress (high acidity, *etc.*) caused stable overexpression of K02112 (F-type H^+^ transporting ATPase). Active primary metabolism of bacteria caused stable overexpression of D-lactate dehydrogenase and glyceraldehyde 3-phosphate dehydrogenase (Dataset 5).

Collectively, our findings from part two of this study supported the second regulation principle, that is, in microbial ecosystems with metabolic division of labour, regulating stability of beneficial distribution for microbiota could help efficiently reduce metabolic instability that caused by biotic factors.

One typical beneficial distribution situation in *baijiu* fermentation is electron acceptors for microbial metabolism. During anaerobic *baijiu* fermentation process, ethanol, lactate and acetate are all electron acceptors with high concentration [68]. During fermentation phase II, we observed similar strong microbial inhibition by lactate, acetate and weak microbial inhibition by ethanol in two unstable batches (Fig. 4b and Fig. 4c). This microbial inhibition pattern could explain why *baijiu* microbiome still transferred raw materials to fermented liquor rather vinegar or other products despite the instability caused by biotic factors. Thus, we could predict fermentation productivity (mainly ethanol) through proper microbial inhibition model to help rationally regulate biotic factors [56]. For other flavor-related metabolic compounds, we need constraint-based metabolic modeling to help ensure stability of key microbial beneficial distribution [30]. Our results presented genes, pathways, and species that enabled future study to improve fermentation stability through flux variability analysis or flux balance analysis [69].

Another challenge that limits fermentation stability is functional redundancy of microbiome. This functional redundancy can provide a level of stability to the fermentation process, as the loss of one microbial species or metabolic pathway can be compensated by the presence of other species or pathways with similar functions [70]. This redundancy also allows for flexibility in the nutrient cycling, as the microbiome can adjust to changes in the environment or substrate availability by different microbial interactions within and among fungal and bacterial communities [71, 72]. However, while functional redundancy can provide stability and resilience to the fermentation microbiome, it can also limit the potential for optimization and improvement of the process. For example, if one microbial species or pathway is more efficient at producing a desired product than others, the presence of redundant pathways may hinder the ability to selectively enhance the desired function [73-75]. *Baijiu* fermentation is a functional redundant ecosystem due to coexistence of microbial species with similar metabolic functions [13]. Our results showed that the stably coexisted core microbial species presented more or less overlapping functional genes, suggesting that partial uncertainty of metabolic outcomes is inevitable due to microbiome functional redundancy. The modernization and standardization for completely stable indigenous liquor fermentation may need microbiome engineering technologies to precisely replace or simplify non-essential redundant microbial activities in time, space and composition [49, 76].

Simplifying complexity of microbial ecosystem are closely associated with biodiversity and stable multiple metabolic functionalities [7, 77]. The overall ecosystem metabolic stability can be increased when biodiversity is low, but decreased when biodiversity is high [7]. In this study, we found that high bacterial diversity and topological complexity of metabolic functional network actually decreased the robustness metabolic network (Fig. 6). This result suggested that the biodiversity of industrial indigenous *baijiu* fermentation has already over the tipping point. We may continue improve industrial fermentation stability by properly decreasing biodiversity [78].

Regulating inoculation ratio is a practical way to change biodiversity. Notably, regulating inoculation ratio changed both fungal and bacterial diversity. In part three of this study, we found decreasing biodiversity by “inoculation ratio” strategy was not effective for optimizing fermentation stability. We argued that decreasing inoculation ratio reduced the fungal contribution to fermentation stability [79]. The “inoculation ratio” strategy may not be effective for optimizing fermentation when fungi and bacteria have opposite contributions to fermentation stability. Nevertheless, we can still enhance fermentation stability by determining the optimal inoculation ratio that maximizes the overall beneficial contributions from both fungi and bacteria.

Regulating inoculation ratio could affect microbial density that may cause different microbial assembly through a dispersal process [80]. In the simulated fermentations, we found raising inoculation ratio of starters (*Daqu*, a sort of Koji) could increase and decrease stochasticity of community assembly (Fig. 7). Notably, high microbial density will accelerate the fermentation through strongly determined assembly pattern that can constrain complexity of metabolic functions [81, 82]. As a result, fermentation quality (flavor complexity, aroma structure, *etc.*) would be harmed [34] despite the high fermentation stability. Thus, our results suggested that we may find the optimal spatial density to optimize both fermentation quality and stability [83]. This finding may help future work to reduce the spatial complexity to establish the fundamental rules of designing the best microbial ratio. Collectively, the findings from part two and part three supported the third regulation principle, that is, in high functional redundancy microbial ecosystems, adjusting biodiversity while ensuring metabolic network complexity of microbiome could help properly reduce metabolic instability that caused by biotic factors.

Overall, we summarized the trinity traits of fermentation microbiome and provided three regulation strategies to stabilize metabolic functions (Fig. 8). The fermentation microbiome is a complex ecosystem that exhibits variable metabolic stability through a combination of redundant metabolic genes, metabolic division of labor between fungi and bacteria, and the modulation of gene expression in response to parameters and environmental cues. Understanding abiotic and biotic factors that drive microbial assembly and the metabolic interactions between microbial species can help regulate microbiome metabolic stability to develop sustainable bioproduction systems.

**Fig. 8.**
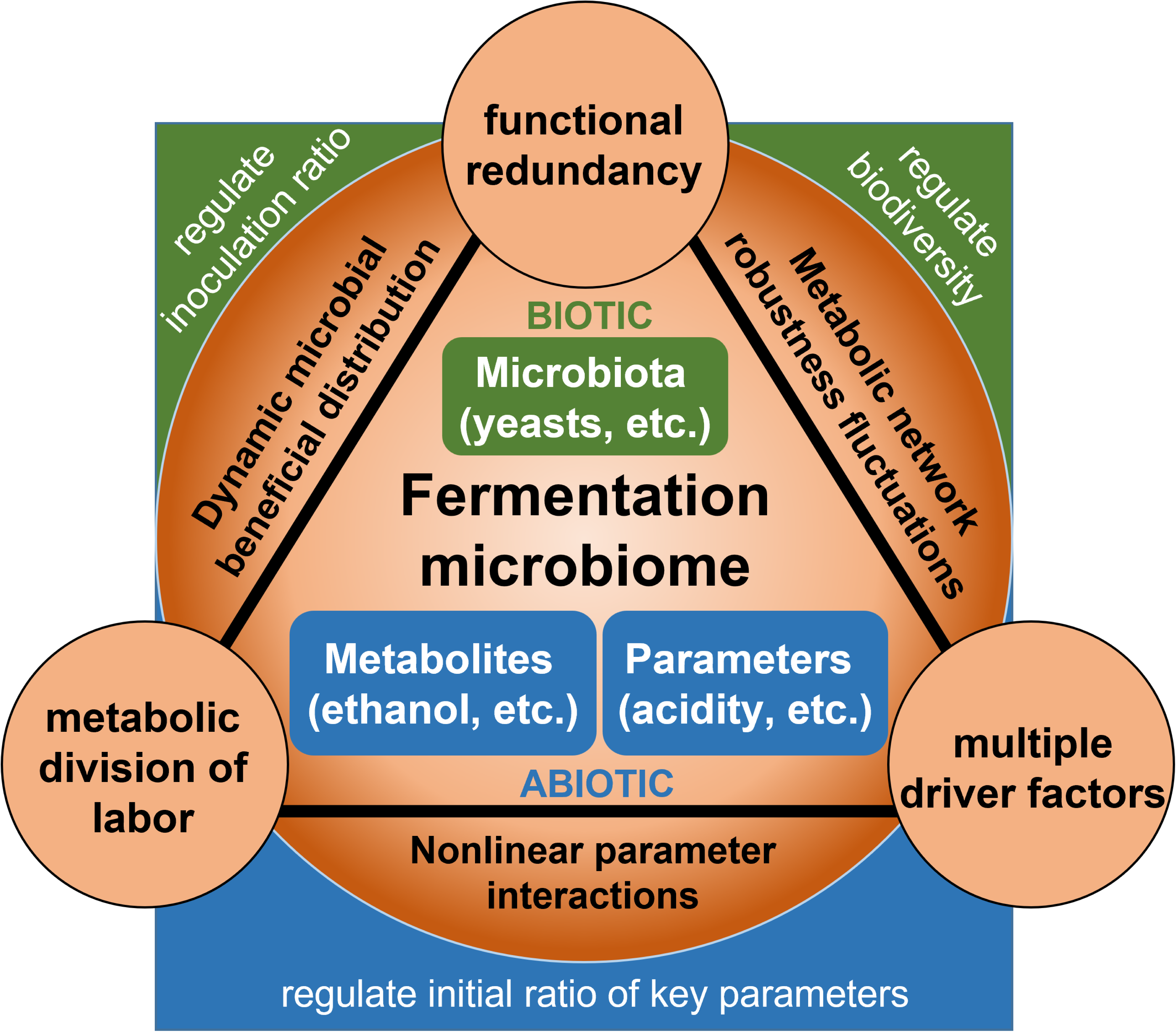
Trinity traits of fermentation microbiome and regulation strategies for stable metabolic functions. Indigenous fermentation microbiome consists of abiotic factors like parameters and microbial metabolites and biotic factors like microbiota. The fermentation microbiome has multiple driving factors of microbial succession. The fermentation microbiome has the metabolic division of labor with efforts from both fungi and bacteria. The fermentation microbiome has redundant metabolic genes to perform metabolic activities. The trinity traits make fermentation microbiome has nonlinear parameters interactions, dynamic microbial beneficial distribution and fluctuate robustness of metabolic network. As a result, fermentation microbiome acts like spheres with variable equilibrium points, representing variable solutions to achieve metabolic objectives. Three regulation strategies act like fixed blocks to stabilize the sphere, representing stability optimization of microbiome metabolism.

## Conclusions

In our three-step experiment, we assessed the effect of biotic and abiotic factors on metabolic functional stability of microbiome in *baijiu* fermentation. Our study demonstrates how, by regulating initial key parameters and initial inoculation ratio, to reduce fermentation instability caused by biotic or abiotic factors accordingly. The stable microbial beneficial distribution and high robustness of metabolic network are essential to metabolic stability of microbiome. Overall, our work introduces theoretical insights and proves the feasibility of enhancing metabolic functional stability through initial conditions in complex indigenous fermentations.

## List of abbreviations

HS-SPME-GC-MS: Headspace Solid-Phase Microextraction Gas Chromatography-Mass Spectrometry
UPLC-MS/MS: Ultra performance liquid chromatography - tandem mass spectrometer
KEGG: Kyoto Encyclopedia of Genes and Genomes
FPKM: Fragments Per Kilobase of transcript per Million fragments
βNTI: β-nearest taxon index
QX: means pure aroma
NX: means pure aroma
JX: means sauce aroma
PCA: Principal Component Analysis
FDR: False discovery rate

## Declarations

### Ethics approval and consent to participate

Not applicable.

## Consent for publication

Not applicable.

## Data availability

Sequencing reads can be accessed in the NCBI database with the accession numbers PRJNA932249 for amplicon sequencing data, PRJNA850152 for metatranscription data, and PRJNA932021 for metagenomic sequencing data.

## Author contribution statement

**Y. T.**: Experiments, Data analysis, Writing - original draft. **Y. Z.**: Writing - review & editing. **R. W.**: Writing - review & editing. **W. S.**: Data analysis, Writing - review & editing. **Y. X.**: Writing - review & editing, Supervision, Funding acquisition. **V. M.**: Writing - review & editing, Supervision. All authors have read and approved the final manuscript.

