## supplementary materials for "Regulating microbiome metabolic stability for stable indigenous liquor fermentation"

**This supplementary file includes:**

Supplementary figures: Fig. S1–Fig. S6

Supplementary tables: Table S1

**Other supplementary information for this manuscript includes:**

Supplementary dataset.xlsx: Dataset 1–Dataset 5

**Dataset 1** Fermentation parameters and productivity types at end of batches.

**Dataset 2** Dynamics of fermentation parameters throughout industrial fermentations.

**Dataset 3** FPKM values of genes in glycolysis, pyruvate metabolism and xenobiotics metabolism.

**Dataset 4** Express level of sub-clusters metabolites.

**Dataset 5** Fungal and bacterial gene overexpression mapping in representative batches

**Supplementary figures:**

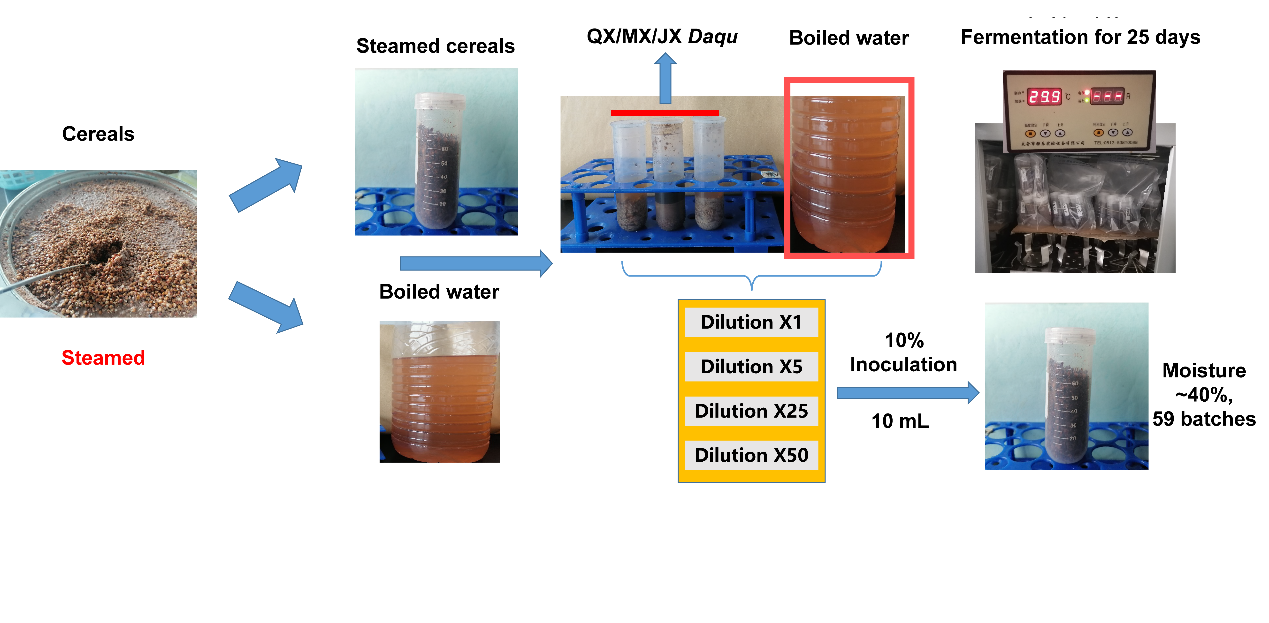

**Figure S1|** Experimental design of simulated fermentations. The external fermentation conditions of all batches were same, including moisture and fermentation temperature. Dilutions of starters were used to adjust initial microbial spatial density.

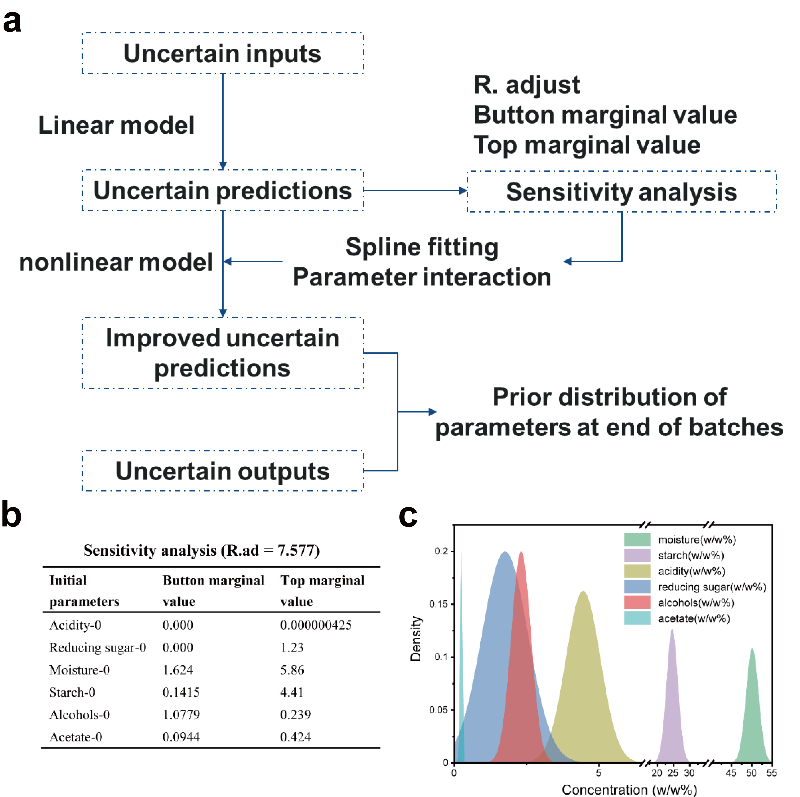

**Figure S2|** Unstable fermentation predictions based on initial parameters. (a) The modelling steps for unstable fermentation predictions. (b) Values of sensitivity analysis (more information see table S1). (c) Prior distribution of parameters at end of batches for potential model optimization (Bayesian inference, *etc.*).
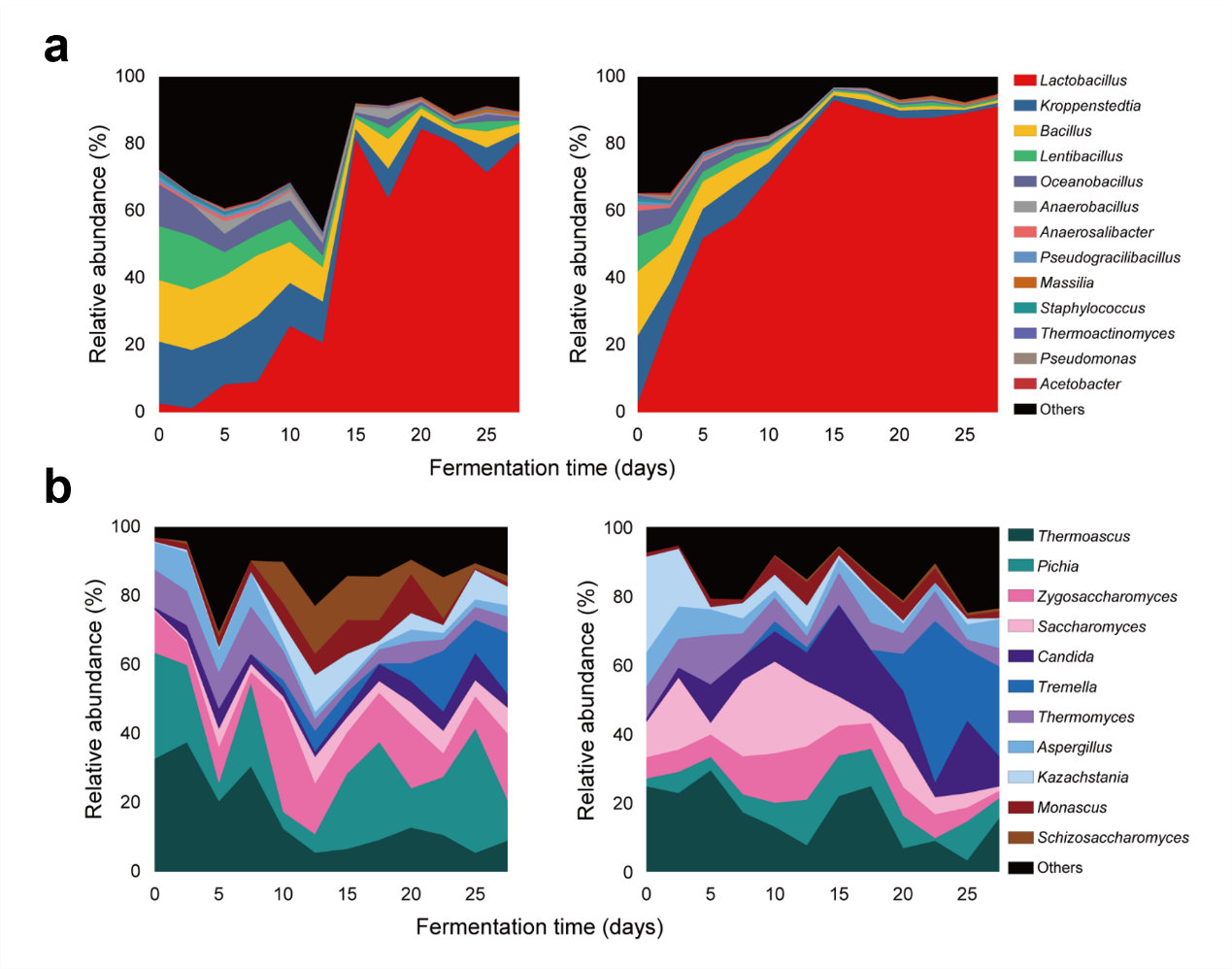

**Figure S3|** (a) Bacterial and (b) fungal succession during fermentation at genera taxonomic level.

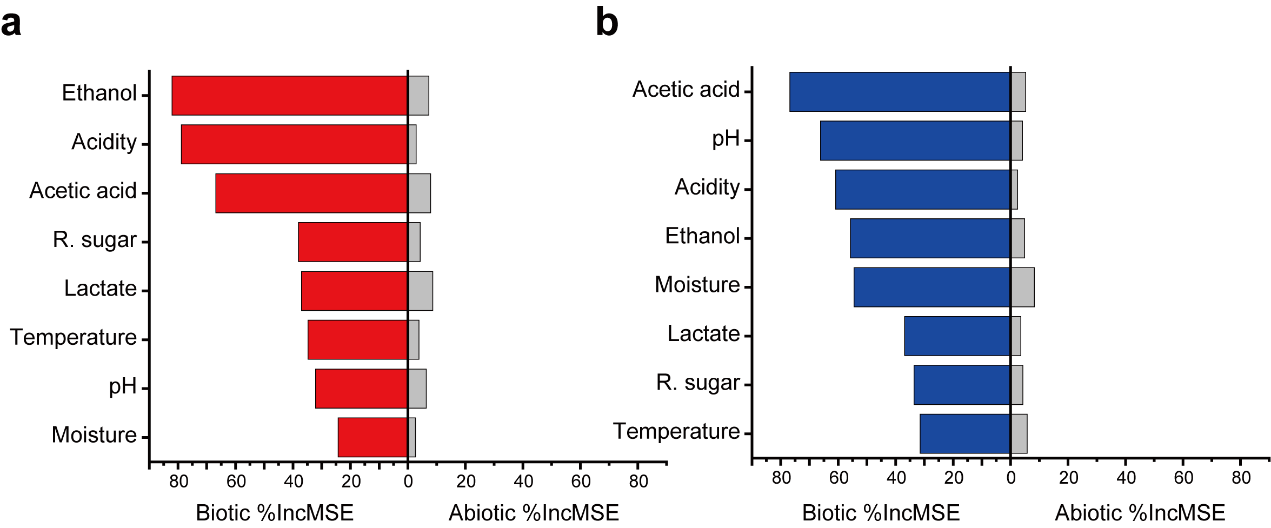

**Figure S4|** Contribution of biotic and abiotic factors to eight fermentation parameters in (a) batch A and (b) batch B by random forest prediction.

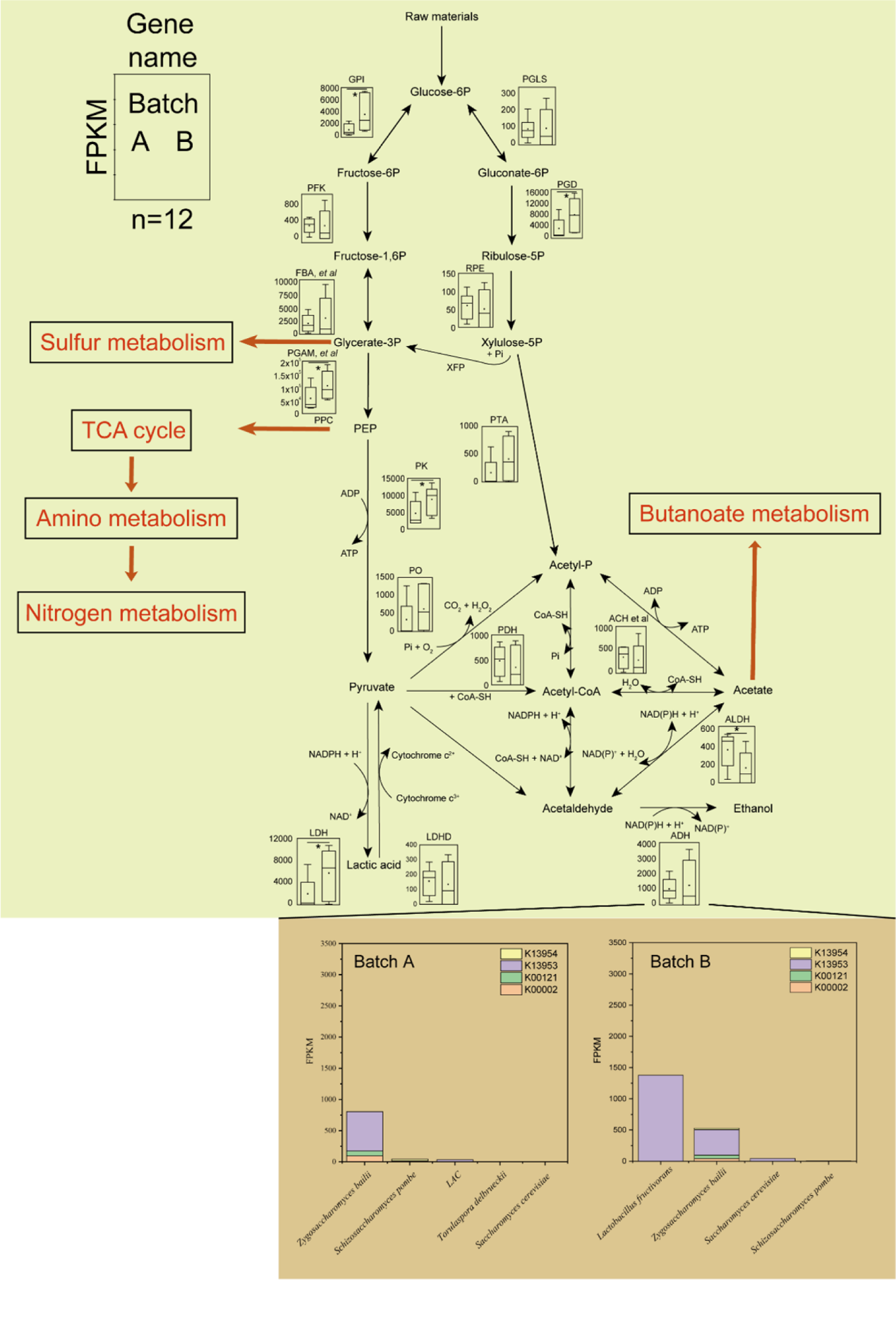

**Figure S5|** Comparison of metabolism flux between two batches of fermentation. Gene transcription difference in primary metabolism. Significance was calculated via pair-sample t test (*P* < 0.05, *). The bar plot in the button shows the alternative core species of ADH expression.

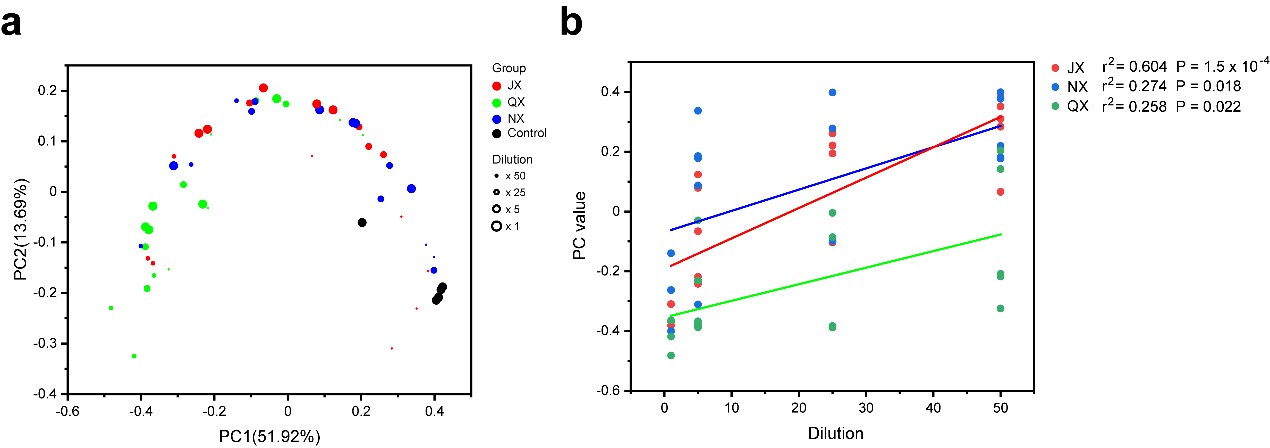

**Figure S6|** Volatile flavor profiles of simulated fermentations. (a) The principal co-ordinates analysis (PCoA) plot of fermentation flavour. (b) Association between dilution and fermentation flavor (one-way ANOVA test).

**Supplementary tables**

**Table S1.** **Sensitivity analysis of linear models**

| **Initial parameters** | **BMV** | | **TMV** | | **Output** |
| --- | --- | --- | --- | --- | --- |
| Acidity-0 | 0.000 | | 0.000000425 | | Alcohols-30(yield) |
| Reducing sugar-0 | 0.000 | | 1.230 | | Alcohols-30(yield) |
| Moisture-0 | 1.624 | | 5.860 | | Alcohols-30(yield) |
| Starch-0 | 0.142 | | 4.410 | | Alcohols-30(yield) |
| Alcohols-0 | 1.078 | | 0.239 | | Alcohols-30(yield) |
| Acetate-0 | 0.094 | | 0.424 | | Alcohols-30(yield) |
| Acidity-0 | | 0.979 | | 1.181 | PC1(quality) |
| Reducing sugar-0 | | 1.751 | | 4.248 | PC1(quality) |
| Moisture-0 | | 0.857 | | 0.000158 | PC1(quality) |
| Starch-0 | | 0.000 | | 0.004599 | PC1(quality) |
| Alcohols-0 | | 7.525 | | 8.337 | PC1(quality) |
| Acetate-0 | | 0.265 | | 0.000184 | PC1(quality) |
| Acidity-0 | | 0.396 | | 1.794 | PC2(quality) |
| Reducing sugar-0 | | 0.046 | | 0.053 | PC2(quality) |
| Moisture-0 | | 0.464 | | 0.088 | PC2(quality) |
| Starch-0 | | 0.854 | | 0.188 | PC2(quality) |
| Alcohols-0 | | 0.291 | | 0.489 | PC2(quality) |
| Acetate-0 | | 0.000 | | 1.165 | PC2(quality) |
| Acidity-0 | | 0.966 | | 2.931 | Acidity-30(quality) |
| Reducing sugar-0 | | 0.000 | | 0.141 | Acidity-30(quality) |
| Moisture-0 | | 0.000 | | 0.021 | Acidity-30(quality) |
| Starch-0 | | 1.039 | | 0.700 | Acidity-30(quality) |
| Alcohols-0 | | 2.250 | | 1.750 | Acidity-30(quality) |
| Acetate-0 | | 0.146 | | 1.115 | Acidity-30(quality) |
| Acidity-0 | | 1.097 | | 1.088 | R. sugar -30(quality) |
| Reducing sugar-0 | | 1.892 | | 4.012 | R. sugar -30(quality) |
| Moisture-0 | | 0.815 | | 0.054 | R. sugar -30(quality) |
| Starch-0 | | 0.136 | | 0.120 | R. sugar -30(quality) |
| Alcohols-0 | | 6.879 | | 8.668 | R. sugar -30(quality) |
| Acetate-0 | | 0.295 | | 0.000346 | R. sugar -30(quality) |
| Acidity-0 | | 0.535 | | 1.097 | Moisture -30(quality) |
| Reducing sugar-0 | | 0.050 | | 4.912 | Moisture -30(quality) |
| Moisture-0 | | 2.809 | | 11.64 | Moisture -30(quality) |
| Starch-0 | | 1.147 | | 11.23 | Moisture -30(quality) |
| Alcohols-0 | | 5.693 | | 0.062 | Moisture -30(quality) |
| Acetate-0 | | 0.676 | | 0.121 | Moisture -30(quality) |
| Acidity-0 | | 0.788 | | 1.186 | Starch -30(quality) |
| Reducing sugar-0 | | 0.000 | | 3.500 | Starch -30(quality) |
| Moisture-0 | | 0.882 | | 6.472 | Starch -30(quality) |
| Starch-0 | | 1.889 | | 7.863 | Starch -30(quality) |
| Alcohols-0 | | 5.594 | | 0.447 | Starch -30(quality) |
| Acetate-0 | | 1.011 | | 0.057 | Starch -30(quality) |
| Acidity-0 | | 0.000 | | 1.656 | Acetate -30(quality) |
| Reducing sugar-0 | | 0.681 | | 0.692 | Acetate -30(quality) |
| Moisture-0 | | 0.666 | | 0.00315 | Acetate -30(quality) |
| Starch-0 | | 1.989 | | 0.911 | Acetate -30(quality) |
| Alcohols-0 | | 0.0413 | | 0.002 | Acetate -30(quality) |
| Acetate-0 | | 0.0315 | | 2.253 | Acetate -30(quality) |
